# BiFET: A <u>Bi</u>as-free Transcription Factor <u>F</u>ootprint <u>E</u>nrichment <u>T</u>est

**DOI:** 10.1101/324277

**Authors:** Ahrim Youn, Eladio J. Marquez, Nathan Lawlor, Michael L. Stitzel, Duygu Ucar

**Affiliations:** The Jackson Laboratory for Genomic Medicine, Farmington, CT, 06032, USA; institute for Systems Genomics, University of Connecticut, Farmington, CT, 06032, USA; Department of Genetics & Genome Sciences, University of Connecticut, Farmington, CT, 06032, USA

## Abstract

Transcription factor (TF) footprinting uncovers putative protein-DNA binding via combined analyses of chromatin accessibility patterns and their underlying TF sequence motifs. TF footprints are frequently used to identify TFs that regulate activities of cell/condition-specific genomic regions (target loci) in comparison to control regions (background loci) using standard enrichment tests. However, there is a strong association between the chromatin accessibility level and the GC content of a locus and the number and types of TF footprints that can be detected at this site. Traditional enrichment tests (e.g., hypergeometric) do not account for this bias and inflate false positive associations. Therefore, we developed a novel method, <u>B</u>ias-free <u>F</u>ootprint <u>E</u>nrichment <u>T</u>est (BiFET), that corrects for the biases arising from the differences in chromatin accessibility levels and GC contents between target and background loci in footprint enrichment analyses. We applied BiFET on TF footprint calls obtained from human EndoC-βH1 ATAC-seq samples using three different algorithms (CENTIPEDE, HINT-BC, and PIQ) and showed BiFET’s ability to increase power and reduce false positive rate when compared to hypergeometric test. Furthermore, we used BiFET to study TF footprints from human PBMC and pancreatic islet ATAC-seq samples to show its utility to identify putative TFs associated with cell-type-specific loci.

## INTRODUCTION

Detecting transcription factor (TF) binding to DNA is critical to understand and study transcriptional control of gene expression (1). Chromatin immunoprecipitation-sequencing (ChIP-seq) assays are effective in uncovering genome-wide binding patterns of a TF. However, profiling multiple TFs using this technology in a cell type of interest is costly and requires large input cell numbers, which limits its wide application to study TF-DNA interactions. A more high-throughput alternative to experimental profiling of these interactions is digital TF footprinting (2), which computationally infers TF binding to DNA by integrating chromatin accessibility patterns (e.g., DNase-seq/ATAC-seq profiles) with the underlying TF binding motifs represented as position weight matrices (PWM) (3,4). Several algorithms have been developed for this purpose to model the probability of a TF’s binding to a given locus from genomewide chromatin accessibility maps (5–8).

Due to advances in genomewide chromatin accessibility profiling, notably the ATAC-seq (9) technology, increasing numbers of chromatin accessibility maps have been generated in primary human cells to study complex diseases, including cancer (10), systemic lupus erythematosus (11), immunosenescence (12,13), and type 2 diabetes (14–16). Effective detection and analyses of TF footprints from these data will be instrumental to nominate potential regulators associated with a clinical phenotype of interest (e.g., immunosenescence (12) or cancer subtypes (17)). TF footprint enrichment analyses can be utilized for this purpose by comparing the number of TF footprint calls in genomic regions of interest (target sites) against footprint calls in a reference set of regions (background sites). Unfortunately, standard enrichment tests (e.g., hypergeometric test or equivalently one-sided Fisher’s exact test) are subject to biases intrinsic to TF footprinting data and can lead to spurious enrichment results unrelated to the biological/clinical question of interest.

In our analyses, TF footprints obtained from ATAC-seq samples in three different human cell/tissue types (EndoC-βH1 pancreatic beta cell line (18), peripheral blood mononuclear cells (PBMCs), and pancreatic islets) revealed two major sources of bias affecting downstream enrichment analyses: differences in sequence GC content and chromatin accessibility levels of target/background regions. First, the GC content of a region significantly affects which TF footprints can be detected in this locus; when target regions on average have higher GC content than the background regions, many GC-rich motifs are falsely identified as enriched in targets, which has been previously noted in motif enrichment analyses and corrected for by minimizing the imbalance of GC content between target and background sites (19,20). TF footprint analyses are subject to a similar bias, however, no current methodology accounts for this bias in TF footprint enrichment analyses.

Second, detection of footprints in an open chromatin region (OCR) is highly dependent on the number of reads (e.g., Tn5 cuts) spanning this region. DNA-cutting enzymes, such as DNase I or Tn5, have sequence-specific biases that contribute to the differences in the number of reads at different OCRs (4,21–23). Footprint detection algorithms typically identify footprints in an OCR using the depletion of cuts at a given sequence relative to nearby flanking regions (24). Therefore, these algorithms likely detect more footprints in OCRs with more cuts (i.e., more read counts). Due to this association between read count numbers at a given locus and the number of footprints detected at this site, standard enrichment tests detect many false positive TFs when target regions have more reads on the average compared to the background regions.

In this study, we present a robust enrichment test for TF footprinting data analyses, BiFET: <u>B</u>ias-free <u>F</u>ootprint <u>E</u>nrichment <u>T</u>est, that corrects for the biases arising from differences between background and target regions in terms of their number of sequencing reads and GC content (**Figure 1**). We applied BiFET on TF footprint calls from EndoC-βH1 ATAC-seq data using three different footprint algorithms: CENTIPEDE (6), HINT-BC (25) and PIQ (7). EndoC footprints from three algorithms were used to simulate true TF binding events, which enabled us to compare the detection power and the false positive rate of BiFET to the frequently used hypergeometric test. In comparison to the hypergeometric test, BiFET is robust to the choice of the background set and has high detection power and low false positive rate regardless of the algorithm used to call footprints. Furthermore, we applied BiFET on ATAC-seq data from human PBMCs and pancreatic islets to uncover TFs that are associated with PBMC or islet-specific regulatory elements and studied the efficacy of BiFET in the downstream enrichment analyses of footprinting data from clinically relevant samples.

**Fig 1.**
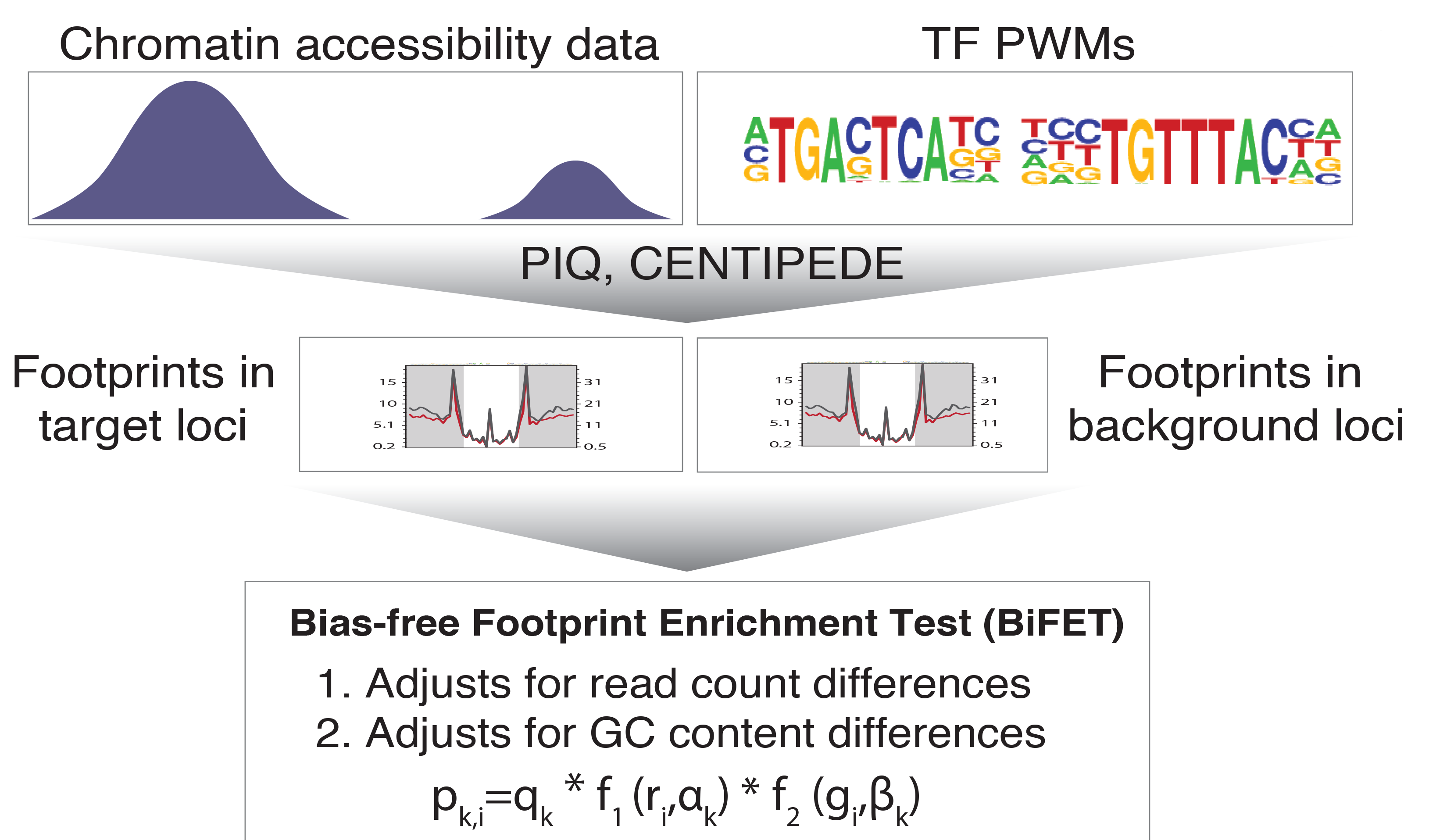
BiFET framework. BiFET models chromatin accessibility (i.e., read count) and GC content differences between target and background regions for an effective TF footprinting enrichment test.

## MATERIAL AND METHODS

### <u>B</u>ias-free <u>F</u>ootprint <u>E</u>nrichment <u>T</u>est (BiFET)

BiFET aims to identify TFs whose footprints are over-represented in target regions (e.g., ATAC-seq peaks associated with a phenotype) compared to background regions after correcting for differences in read counts and GC content between target and background regions. Specifically, BiFET tests the null hypothesis that target regions have the same probability of having footprints for a given TF *k* as the background regions after correcting for the read count and the GC content bias (See **Figure 1** for a summary of the proposed framework). For this, the number of target peaks with footprints for TF *k* (*t*_*k*_) is used as a test statistic and the p-value is calculated as the probability of observing *t*_*k*_ or more peaks with footprints under the null hypothesis. The association between read counts and footprint detection rate, is modeled with a logistic function *f*_1_:

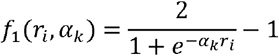

where *r*_*i*_ denotes the number of reads in peak *i*. *f*_1_ is equal to 0 when the peak has no reads (*r*_*i*_ =0) and increases monotonically converging to 1 as the number of reads increases to infinity at a rate determined by *α*_*k*_ >0 (See Supplementary **Figure S1A** for the relation between *f*_1_ and *α*_*k*_ for increasing read count values).

Similarly, we model the association between the GC content of a genomic region and the footprint detection by introducing a second logistic function *f*_2_:

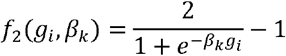

where *g*_*i*_ denotes the GC content (proportion of GC) in the genomic region *i* and *β*_*k*_ > 0 determines how fast *f*_2_ converges to 1. Unlike the read count bias, the positive association between the GC content and footprint detection exists only for TFs with GC-rich motifs (See **Figures 2C** and **D** and **Supplementary Figures 2B, C, E,** and **F** for the relation between footprint detection and GC content of genomic regions for GC-rich and GC-poor motifs). The logistic function *f*_2_ with various values of *β*_*k*_ can model this TF-specific association between GC content and the footprint detection. For example, when *β*_*k*_ is high (i.e., 10,000) as in **Supplementary Figure 1B**, *f*_2_ is equal to 1 for any value of *g*_*i*_ >0, hence the footprint detection does not depend on the GC content. For GC-poor motifs, *β*_*k*_ will have a high value, hence there will not be an association between the GC content and the footprint detection.

Finally, the probability that a footprint for TF *k* is called in a peak *i* (*p*_*k,i*_) is modeled as:

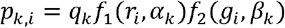

In this model, the parameter *q*_*k*_ denotes TF-specific binding rate, which is adjusted by functions *f*_1_ and *f*_2_ that measure the effect of the read count levels and GC content of peaks on footprint detection rates. This model assumes that the read counts and the GC contents of genomic regions independently affect the probability of footprint detection. This assumption is supported by our analyses (**Supplementary Figure S3**), which shows that the relation between footprint detection and GC content (or read counts) is preserved as we stratify the data by read counts (or GC content), respectively.

When target and background regions have similar read counts and GC content, the difference in rates of TF footprint calls can be explained by the difference in *q*_*k*_ between the two sets. Therefore, we test if *q*_*k*_ differs between the target and background regions. More specifically, we assume that the probability of the target peak *i* having a footprint for TF *k* is *p*_*k,i,1*_ = *q*_*k,1*_*f*_1_(*r*_*i*_,*α*_*k*_)*f*_2_(*g*_*i*_,*β*_*k*_) and the probability of the background peak *i* having a footprint for TF *k* is *p*_*k,i,2*_ = *q*_*k,2*_*f*_1_(*r*_*i*_,*α*_*k*_)*f*_2_(*g*_*i*_,*β*_*k*_) and test the null hypothesis (*H*_o_: *q*_*k,1*_ = *q*_*k,2*_ = *q*_*k*_) and estimate the parameters *q*_*k*_,α_*k*_ and *β*_*k*_ by maximizing the likelihood of the footprint data for TF *k*:

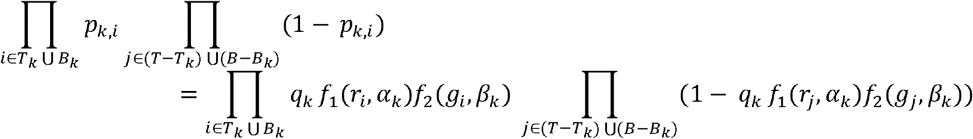

where *T* and *B* denote target and background peaks and *T*_*k*_ and *B*_*k*_ are target peaks and background peaks with footprints for TF *k*, where |*T*_*k*_| = *t*_*k*_ and |*B*_*k*_| = *b*_*k*_. The optimization was performed by R optim function with a limited-memory modification of the BFGS quasi-Newton method (26).

We then define the p-value for testing the null hypothesis as the probability that there are *t*_*k*_ or more target peaks with footprints for TF *k*:

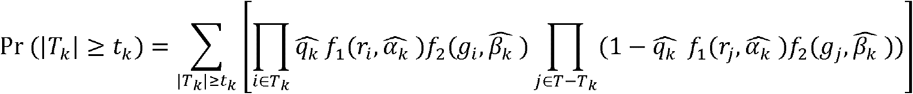

where 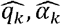 and 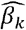 are maximum likelihood estimates (MLE) of *q*_*k*_,*α*_*k*_ and *β*_*k*_. This probability is calculated using R package poibin (27).

BiFET is available as a Bioconductor package named “BiFET”. Instructions on how to use BiFET and the required input files are available at https://github.com/UcarLab/BiFET/blob/master/vignettes/BiFET.Rmd.

### Simulation studies in EndoC cell line

#### 1. EndoC ATAC-seq data processing

We assessed the performance of BiFET by simulating TF footprint calls using ATAC-seq data in human EndoC-ßH1 beta cell line (18). From these data 138,707 OCRs (i.e., ATAC-seq peaks) were identified using *MACS* version 2.1.0 (28) with parameters “-nomodel-f BAMPE”. The peaks were truncated to a total length of 200bp (+/− 100bp from the peak center) to eliminate biases associated with differences in peak lengths. This same peak length cut-off has been used in all of our analyses to ensure that number of footprints was not affected by differences in target and background peak lengths.

#### 2. Footprint calling using three algorithms

A number of footprinting algorithms have been developed to predict TF binding sites using DNase-seq or ATAC-seq data, which broadly fall into two categories: shape detection and motif-driven. Shape detection algorithms, e.g. Neph (29), Wellington (30), DNase2TF (8), Boyle (31), HINT (25), and HINT-BC (32) scan DNase-seq or ATAC-seq data to detect a footprint-like spatial shape—short genomic regions of low (DNase I or Tn5) cleavage immediately flanked from both ends by high cleavage— without specifying the TF motif. Motif-driven algorithms on the other hand, e.g., FLR (33), CENTIPEDE (6), PIQ (7), and BinDNase (34), first scan the genome for known TF sequence motifs and classify loci with a motif as bound or unbound based on the chromatin accessibility profiles (32). To evaluate BiFET’s performance for different TF footprinting detection methods, we chose three frequently used algorithms: HINT-BC (representing shape detection algorithms), CENTIPEDE, and PIQ (representing motif-driven algorithms) to call TF footprints from EndoC-ßH1 ATAC-seq data.

CENTIPEDE uses a Bayesian mixture model to estimate the posterior probabilities of each motif site bound by the corresponding TF (6). On the other hand, PIQ uses a Gaussian process to model and smooth the footprint profiles around motif sites to estimate the probability of occupancy for each motif occurrence (7). HINT-BC (HINT bias-corrected) is an extension of the method HINT (<u>H</u>mm-based <u>I</u>de<u>N</u>tification of <u>T</u>f footprints), which adjusts for the sequence cleavage bias of cutting enzymes used in chromatin accessibility assays (32).

We applied all three algorithms with their default parameters using a PWM library compiled from the JASPAR database (35) and Jolma *et al.* (36) (n= 979 PWMs in total). Since HINT-BC does not specify which TF is associated with the detected footprint, we overlapped HINT-BC footprints with this PWM library. In this analysis, if at least 2/3 of a TF’s motif overlapped with a HINT-BC footprint, we associated this TF to the footprint. For all three algorithms TF footprints were filtered based on the scores that measure the confidence of the footprint detection, i.e., positive predictive values (PPV) > 0.9 for PIQ, posterior probabilities of binding > 0.95 for CENTIPEDE and tag-count score > 80^th^ percentile for HINT-C with frequently used thresholds.

#### 3. TF footprinting simulations

To investigate the impact of read count and GC content differences between target and background regions on the enrichment test results, we applied three different methods to select target regions comprising 5% of all EndoC ATAC-seq peaks (6,935 peaks):

1. Target peaks were randomly selected from all peaks (target + background) so that the expected read counts and GC content do not differ between target and background regions.
2. Target peaks were randomly selected by setting the sampling probability to be proportional to f(x=read counts per peak) using four functions: (a) f(x)=x, (b) f(x)=x^1/2^, (c) f(x)=x^−1/2^, and (d) f(x)=x^−1^ where the average read count for target peaks decreases from (a) to (d). In (a) and (b), target peaks have higher read counts than the background peaks, whereas in (c) and (d), they have lower read counts than the background peaks.
3. Target peaks were randomly selected by setting the sampling probability to be proportional to f(x= GC content per peak) using four different f functions: (a) f(x)=x, (b) f(x)=x^1/2^, (c) f(x)=x^−1/2^, (d) f(x)=x^−1^ where the average GC content for the target peak set decreases from (a) to (d). In (a) and (b), the average GC content for target peaks are higher than that of background peaks, whereas in (c) and (d), it is lower than the background peaks.

In all three cases, target peaks were randomly selected independent of their location, functional association, or TF motif enrichments. Therefore, no TFs were expected to specifically bind to these random peaks, and any TF that is significantly enriched in target peaks is marked as a false positive call.

To quantify the detection power of our method, we randomly selected 10 TFs; for each of these TFs, we simulated artificial footprint calls in N% of the target sets. In other words, for each selected TF *k*, we increased the number of target peaks with footprints for this TF (i.e., |*T*_*k*_|) by N%. We set N to be the binding rate of the TF (i.e., the percentage of peaks with footprints for the TF) across all peaks or across target peaks, whichever is larger. Since we simulated additional footprints for these 10 TFs only within target regions, they should be truly enriched in target peaks compared to the background peaks. Hence, these 10 TFs are treated as true positives (TP) in our analyses, whereas the rest of the TFs detected are considered false positives (FP). Each simulation setting was repeated 50 times to eliminate biases stemming from random samplings. For each simulation, we identified TFs that are enriched in the target set compared to the background set using hypergeometric test and BiFET and assessed the false positive rate and true positive rate for each method using TF footprints from three different footprint detection algorithms.

### Analysis of human islet and PBMC ATAC-seq data

#### 1. Islet and PBMC ATAC-seq data processing

ATAC-seq peaks from five human PBMCs (12) and five human islets (14,16) were called using *MACS* version 2.1.0 with parameters “-nomodel-f BAMPE”. The peaks from all ten samples were merged to generate one consensus peak set (N = 57,108 peaks) by using R package *DiffBind_2.2.5.* (37), where only the peaks called at least twice (out of 10 samples) were included in the analysis. We used the “summits” option to re-center each peak around the point of greatest read overlap and obtained consensus peaks of same width (200 bp, +/− 100bp around the summit). Out of these consensus peaks, we defined PBMC-specific peaks as those that were called in at least four PBMC samples and in none of the islet samples (n=4106 peaks). Similarly, we defined islet-specific peaks as those called in at least four islet samples but in none of the PBMC samples (n=12886 peaks). Consensus peaks that exclude PBMC/islet-specific peaks were used as the background (i.e., non-specific) regions in our enrichment analyses (n=40116 peaks). PIQ was used to call TF footprints from the pooled islet and pooled PBMC samples to increase the detection power for TF footprints based on JASPAR PWMs (n=454 in total). Only the TF footprints with positive predictive values greater than 0.9 are used in downstream enrichment analyses.

#### 2. Footprinting calls using random motifs

Unlike in our simulation study, in real world datasets we typically do not know which TFs are true or false positive regulators of the loci of interest. To quantify BiFET’s ability to reduce false positive rates, we generated artificial PWMs and used PIQ to call footprints for these artificial motifs in ATAC-seq samples (i.e., false positive calls). To generate artificial PWMs, we started with the JASPAR PWMs (n=454) and randomly permuted every column (base pair) of the PWM matrix to obtain a random PWM matrix. For each randomly generated PWM (454 in total), we calculated its Euclidean distance to the JASPAR PWMs using R package PWMsimilarity (38) and selected the top 200 random motifs that are the most dissimilar to the known motifs based on their PWM similarity. These 200 random motifs were used to call PIQ footprints from islet and PBMC ATAC-seq samples and used for assessing false positive rates.

## RESULTS

### Number of ATAC-seq reads and GC content of a region affect TF footprints detected at this locus

From EndoC-ßH1 ATAC-seq data, 15,219,923 significant CENTIPEDE footprints were detected for 793 (out of 979 tested) PWMs (Methods). Only 974,975 (6.4%) of these overlapped ATAC-seq peaks that mapped to 790 distinct PWMs. PIQ detected 5,057,304 significant footprints for 978 TF motif PWMs, where 830,795 (16.4%) footprints for 969 PWMs overlapped EndoC ATAC-seq peaks. On the other hand, by design, HINT-BC detects footprints within a given set of regions. In total, 135,657 footprints were detected by HINT-BC that was associated with 979 PWMs (**Figure 2A**). Only the footprints that are within ATAC-seq peaks were used in downstream analysis.

**Fig 2.**
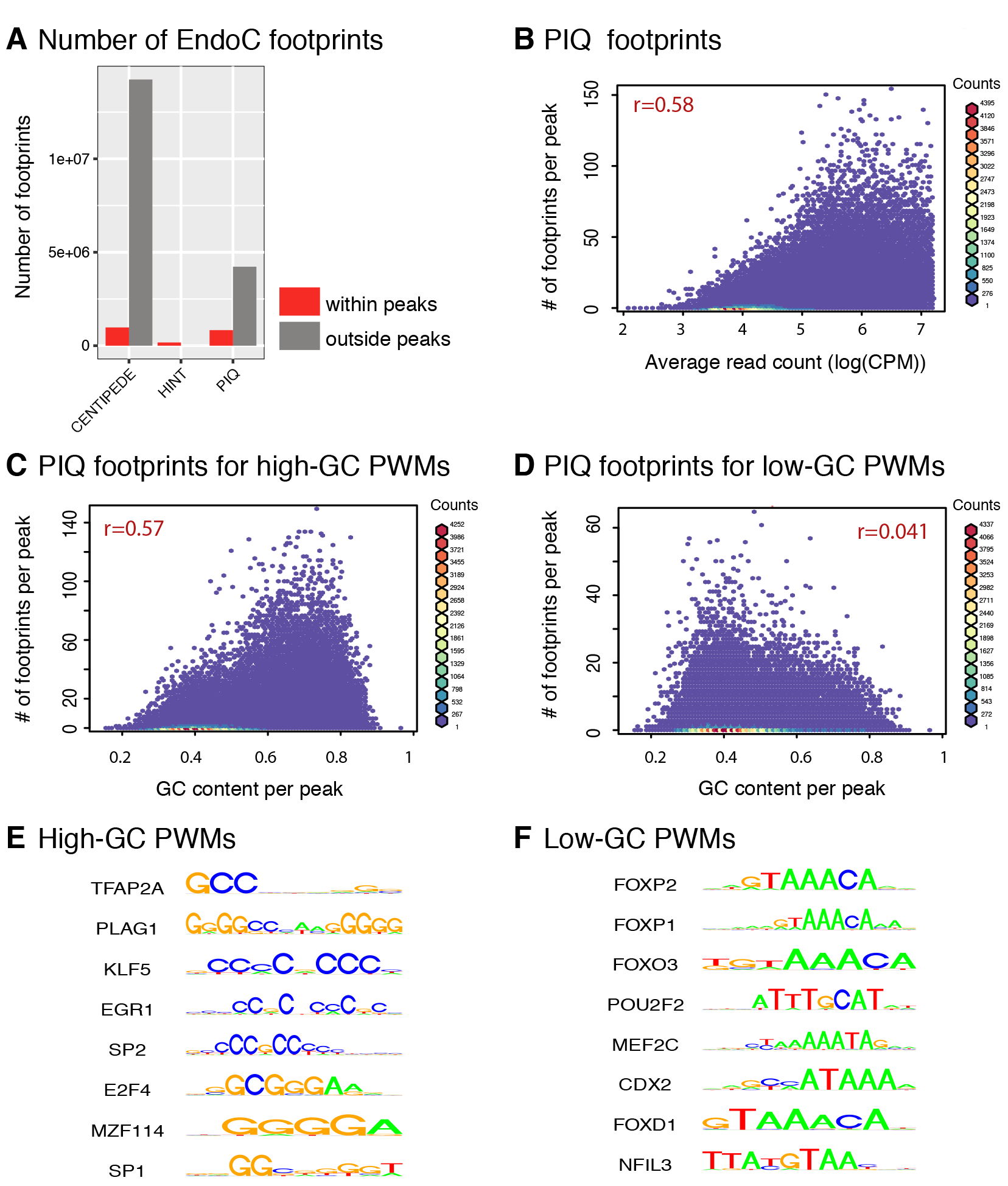
The relation between TF footprints and sequence/genomic features of a locus. (**A**) Number of CENTIPEDE, HINT-BC and PIQ footprints detected in EndoC cell line within (red bars) and outside (gray bars) EndoC ATAC-seq peaks (**B**) ATAC-seq read counts vs. number of PIQ footprints detected in a peak. Due to the outliers, we restricted analyses to peaks whose read counts are below the 99^th^ percentile. (**C**) For TFs with high-GC motifs, GC content of a peak correlate significantly with the number of PIQ footprints detected at this peak. (**D**) For TFs with low-GC motifs, GC content of a peak is not correlated with the number of PIQ footprints detected at this peak. (**E**) Example high-GC content PWMs (**F**) Example low-GC content PWMs.

Despite the differences in genome-wide footprint calls, comparable numbers of footprints were detected within ATAC-seq peaks per TF using different algorithms (Pearson correlation coefficient r=0.58 for CENTIPEDE and PIQ, r=0.72 for HINT-BC and PIQ, r=0.46 for CENTIPEDE and HINT-BC; **Supplementary Figures S4A, B, C**). Furthermore, similar numbers of footprints were detected per peak by different methods (r=0.6 for CENTIPEDE and PIQ, r=0.42 for HINT-BC and PIQ, r=0.32 for CENTIPEDE and HINT-BC; **Supplementary Figures S4D, E, F**), suggesting that different algorithms produce comparable footprints from the same data and they are subject to similar biases in footprint calls.

The number of ATAC-seq reads spanning a peak correlated significantly (p<e−16) with the number of footprints detected within this peak, for all three algorithms: r=0.58 for PIQ (**Figure 2B**), r=0.38 for CENTIPEDE (**Supplementary Figure S2A**), and r=0.35 for HINT-BC (**Supplementary Figure S2D**). Furthermore, for GC-rich motifs (i.e., motifs for which the average probability of having G or C in their PWM matrix > 0.5 such as KLF5 and SP1 in **Figure 2E**), GC content of the peak and the number of footprints detected from this region was also significantly correlated: r=0.57 for PIQ (**Figure 2C**), r=0.54 for CENTIPEDE (**Supplementary Figure S2C**), and r=0.24 for HINT-BC (**Supplementary Figure S2F**). We observed that HINT-BC is less subject to such GC bias, likely because it is not motif-driven and it adjusts for the sequence cleavage bias of cutting enzymes. For TFs with low-GC content PWMs (e.g., Forkhead (FOX) transcription factor family members, POU2F2 in **Figure 2F**), GC content of the peak is not associated with the number of footprints detected at the peak (**Figure 2D**, and **Supplementary Figures S2B, E**). These observations suggest a relationship between locus-specific read count and GC content and the detection probability of TF footprints from this site, which is conserved across three algorithms and likely bias downstream enrichment analyses.

### BiFET enrichment results are robust to differences between target and background regions

By simulating TF footprint enrichments in EndoC cells, we quantified the impact of enrichment test choice under different scenarios (Methods). First, we observed that, as expected, BiFET and hypergeometric test (HT) performs similarly when target and background regions have comparable read counts and GC contents (**Table 1A** for PIQ, **Supplementary Table S1A** for CENTIPEDE and **Supplementary Table S2A** for HINT-BC results).

**Table 1:** Simulation results for EndoC PIQ footprints shows efficacy of BiFET. We calculated the median read counts and GC proportions of target and background sets and the number of true positives (TP), true positive rate (TPR), number of false positives (FP) and false positive rate (FPR) under FDR 0.05 averaged across 50 simulations for each simulation setting: (**A**) randomly sampling target peaks among all peaks, (**B**) randomly sampling target peaks with different read counts among all peaks, and (**C** randomly sampling target peaks with different GC contents among all peaks.

| Median reads<br>target | Median reads<br>background | Median GC %<br>target | Median GC%<br>background | TP (TPR)<br>BiFET | TP (TPR)<br>HT | FP (FPR)<br>BiFET | FP (FPR)<br>HT |
| --- | --- | --- | --- | --- | --- | --- | --- |
| 72 | 72 | 0.43 | 0.43 | 8 (0.8) | 8.1 (0.81) | 0.14 (0.00015) | 0.28 (0.00029) |

B. Randomly sampling target peaks with different read counts
| Simulation<br>setting | Median reads<br>target | Median reads<br>background | TP (TPR)<br>BiFET | TP (TPR)<br>HT | FP (FPR)<br>BiFET | FP (FPR)<br>HT |
| --- | --- | --- | --- | --- | --- | --- |
| a | 275 | 70 | 9.2 ( 0.92 ) | 10 ( 1 ) | 1.3 ( 0.0014 ) | 648 ( 0.68 ) |
| b | 123 | 71 | 8.7 ( 0.87 ) | 9.9 ( 0.99 ) | 0.86 ( 9e-04 ) | 423 ( 0.44 ) |
| c | 58 | 73 | 8.4 (0.84) | 6 (0.6) | 0.06 (6.3e-05) | 0 (0) |
| d | 52 | 74 | 8.7 (0.87) | 5 (0.5) | 0.04 (4.2e-05) | 0 (0) |

C. Randomly sampling target peaks with different GC contents
| Simulation<br>setting | Median GC %<br>target | Median GC %<br>background | TP (TPR)<br>BiFET | TP (TPR)<br>HT | FP (FPR)<br>BiFET | FP (FPR)<br>HT |
| --- | --- | --- | --- | --- | --- | --- |
| a | 0.45 | 0.42 | 8.4 ( 0.84 ) | 9.4 ( 0.94 ) | 22 ( 0.023 ) | 128 ( 0.13 ) |
| b | 0.44 | 0.42 | 8.1 ( 0.81 ) | 8.9 ( 0.89 ) | 0.94 ( 0.00098 ) | 30 ( 0.032 ) |
| c | 0.42 | 0.43 | 8.3 (0.83) | 8 (0.8) | 0.35 (0.00036) | 0.29 (3e-04) |
| d | 0.41 | 0.43 | 8.2 (0.82) | 7.6 (0.76) | 0.61 (0.00064) | 0.24 (0.00026) |

However, when target regions harbor more ATAC-seq reads (i.e., higher read counts) compared to background regions, HT produces large numbers of false positive enrichments. For example, HT identified 648 out of 959 TF motifs (i.e., 969 PWMs detected within peaks - 10 true positives) to be significantly enriched in randomly selected target regions (False positive rate (FPR) = 68%) when there is a significant difference between target and background regions in terms of median read counts (**Table 1B**, setting a). For the same scenario, BiFET controlled the false positive rate at 0.001, where only 1 out of 959 TF motifs had a significant enrichment. On the contrary, when read counts of target regions were lower than those of background regions, HT had a lower True Positive Rate (TPR) than BiFET (e.g., 87% TPR with BiFET vs. 50% with HT for setting d in **Table 1B**). BiFET and HT generated similar results for footprints called using CENTIPEDE and HINT-BC (**Supplementary Table S2B, S3B**)

BiFET also outperformed HT under varying GC content distributions for background and target regions. When the median GC content of target regions is higher than that of the background regions, HT produced many FP calls. For example, 128/959 TF motifs tested (FPR=13%) were detected to be significantly enriched when GC contents of background and target regions were significantly different (**Table 1C**, setting a). Under the same scenario, BiFET better controlled the false positive rate and detected only 22 TFs to be enriched out of 959 (FPR=2%). Similarly, BiFET outperformed HT for footprints obtained from CENTIPEDE (**Supplementary Table S1C**) and HINT-BC (**Supplementary Table S2C**). These simulation results suggest that in comparison to the standard enrichment test (i.e., hypergeometric test), BiFET is robust to the choice of background regions and has high detection power and low false positive rate irrespective of the algorithm used for footprinting calls.

### BiFET uncovers TFs associated with cell-specific regulatory elements

We used BiFET to detect TFs associated with cell-specific OCRs by comparing ATAC-seq data from human PBMCs (12) and pancreatic islets (14). Using a stringent definition of cell-specific accessibility (Methods), we identified 4,106 PBMC-specific ATAC-seq peaks (e.g., *CD28* locus in **Figure 3A**) and 12,886 islet-specific ATAC-seq peaks (e.g., *ISL1* locus in **Figure 3B**). The remaining ATAC-seq peaks (n=40,116) were considered non-specific and used as the background set in our enrichment analyses. PIQ detected 389,948 significant footprints for 401 PWMs within PBMC ATAC-seq peaks and 390,502 significant footprints for 414 PWMs within islet ATAC-seq peaks. Using BiFET and HT, we identified PWMs whose footprints were enriched in PBMC-specific peaks compared to the background peaks (i.e., non-specific peaks) and, similarly, TFs whose footprints were enriched in islet-specific peaks compared to the background peaks. PBMC-specific peaks (i.e., target peaks) had higher ATAC-seq read counts than the background peaks in the PBMC samples, where median log read count of target peaks was 4.8 and median log read count of background peaks was 3.8 (**Figure 3C**, left panel). On the other hand, PBMC-specific peaks had lower GC content than the common peaks (median GC proportion=0.495 vs. 0.53; **Figure 3C**, right panel). Since background peaks had significantly lower read counts than the target peaks, they tended to have fewer footprints. Therefore, if read count bias was not adjusted for, the standard enrichment tests would identify many false positive enrichments.

**Fig.3.**
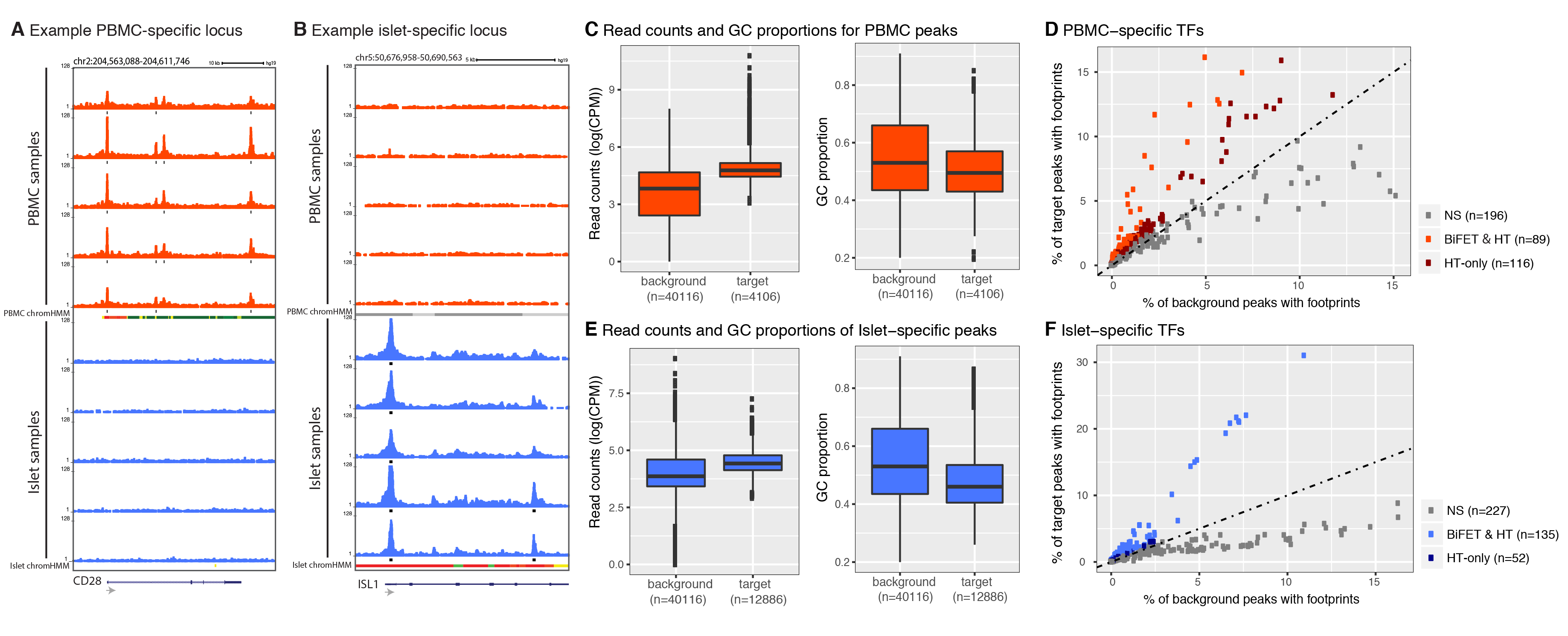
Footprints enriched in PBMC and islet-specific ATAC-seq peaks. (**A**) UCSC genome browser track for example PBMC-specific peaks located around the *CD28* locus. Chromatin accessibility maps in PBMCs (islets) are shown in red (blue). ChromHMM (44) states for PBMCs and islets are represented as colored bars. (**B**) Example islet-specific peak located around the promoter of *ISL1.* (**C**) Read counts (left panel) and GC content (right panel) of PBMC-specific (target) peaks vs. background peaks in PBMC samples. (**D**) Percent of target peaks with footprints vs. percent of background peaks with footprints for each TF in PBMC samples. The TFs that are significant by both BiFET and hypergeometric test are labeled ‘BiFET & HT’ and indicated by red dots, those that are significant only by the hypergeometric test (‘HT-only’) are colored in dark red, and the TFs that are not significant (‘NS’) by either method are colored in gray. (**E**) Read counts (left panel) and GC contents (right panel) of islet-specific (target) peaks vs. background peaks in islet samples. (**F**) Percent of target peaks with footprints vs. percent of background peaks with footprints for each TF in islet samples. The TFs that are significant by both BiFET and hypergeometric test are labeled ‘BiFET & HT’ and colored in blue, those that are significant only by the hypergeometric test (‘HT-only’) are colored in dark blue and the TFs that are not significant (‘NS’) by either method are colored in gray.

BiFET identified 89 PWMs (mapping to 84 TFs) to be significantly (FDR ≤5%) enriched in PBMC-specific peaks out of 401 PWMs that were tested. In comparison, HT identified 205 PWMs as significantly enriched in PBMC-specific peaks, including all 89 PWMs captured by BiFET. As expected, when a PWM is significantly enriched by either method, the percent of target peaks with footprints is higher than the percent of background peaks with footprints for this TF (**Figure 3D**, red dots). However, differences in percent of peaks with footprints between target and background were smaller for the TFs that are solely identified by HT (i.e., dark red dots labeled as ‘HT-only’ in **Figure 3D**).

Similarly, we identified TF footprints enriched in islet-specific peaks using BiFET and HT. Similar to the PBMC data, islet-specific peaks (target peaks) had higher average ATAC-seq read count than the background peaks in islets, where median log read count for target peaks is 4.4 and median log read count for background peaks is 3.9 (**Figure 3E**, left panel). Islet-specific peaks also had lower GC content than the background peaks (median GC proportion = 0.46 vs. 0.53; **Figure 3E**, right panel). BiFET identified 135 PWMs (mapping to 122 TFs) out of 414 tested to be significantly enriched in islet-specific peaks (FDR=0.05), while HT identified 187 PWMs, including the 135 PWMs detected by BIFET. We noted that since the difference in read counts between target and background peaks was not as striking as in PBMC samples (Figures 3C vs. 3E), the number of PWMs exclusively detected using HT were less in islet samples compared to PBMC samples (52 vs. 116). As expected, TFs enriched in islet-specific peaks had more footprints in target regions than in background regions (**Figure 3F**). TFs with significant enrichment according to both methods (light blue dots in **Figure 3F**) clearly separated from the nonsignificant TFs, while the TFs identified only by the HT (dark blue dots in **Figure 3F**) had similar footprint rates between background and target sets, suggesting that enrichments detected only by HT are likely false positives.

To study the functional relevance of TF enrichments obtained from PBMC-and islet-specific peaks, we performed pathway enrichment analysis using HOMER (19). Of the 84 PBMC-specific TFs and 122 islet-specific TFs (Supplementary Table S3) identified by BiFET, 46 TFs were common (**Supplementary Figure S5A**) suggesting that some TFs that regulate cell-specific regions can be common across cell types. The top 3 enriched Wiki pathways for PBMC-specific TFs (n=38) were all immune-related including “Type II, III interferon signaling” and “Development of pulmonary dendritic cells and macrophage subsets” (**Supplementary Table S4**). In contrast, islet-specific TFs (n=76) included *HNF1A, HNF1B, HNF4A,* and *PAX6* (**Supplementary Table S5**), and the most enriched KEGG pathway for islets was “Maturity Onset Diabetes of the Young”. These functional enrichment results show that islet/PBMC-specific TFs identified by BiFET reflect functional enrichments relevant to the cognate cell type.

We repeated the pathway enrichment analyses for TFs identified by HT. HT identified 175 PBMC-specific TFs and 167 islet-specific TFs, of which 113 were common between two cell types (**Supplementary Figure S5B**). We found that the pathways enriched for TFs that are PBMC-specific (n=62) included immune-related pathways, but their p-values were less significant compared to those obtained from BiFET results (**Supplementary Figures S5C, E**; **Supplementary Table S6**). Likewise, we observed that pathways enriched for islet-specific TFs (n=54) had less significant p-values compared to BiFET results for islet biology related pathways (**Supplementary Figures S5D, F**; **Supplementary Table S7**). These results indicate that BiFET was more effective in detecting cell type-specific regulators than the standard enrichment test and can be effective in reducing false positive enrichments between TFs and genomic regions of interest to study human diseases and biology.

### BiFET reduces false positive associations in ATAC-seq footprinting analyses

Although pathway enrichment analysis suggested that the TFs identified by BiFET better capture regulators of PBMC/islet-specific functions, it is difficult to assess which of these are true regulators in clinical samples. To demonstrate the advantage of BiFET in reducing false positives in clinically relevant comparisons, we performed enrichment analyses using BiFET and HT on PIQ footprints for 200 artificially generated random motifs (Methods). For these artificial motifs, 121,085 footprints were detected within PBMC ATAC-seq peaks, where 194 motifs had at least one footprint. The number of detected footprints for these random motifs was highly correlated (Pearson correlation r=0.71) with the read counts similar to the original JASPAR motifs (r=0.66) (**Supplementary Figures S6A, B**) Application of BiFET on these footprints identified 12 PWMs that are significantly enriched in PBMC-specific peaks compared to background peaks, while HT identified 79 significantly enriched PWMs for the same analyses, including all 12 PWMs captured by BiFET. For these random PWMs, the percent of target peaks with footprints was overall lower than that of the original JASPAR motifs (**Supplementary Figure S6C** vs. **Figure 3D**). As expected, for significantly enriched PWMs, percent of target peaks with footprints was higher than the percent of background peaks with footprints (**Supplementary Figure S6C** red dots). Similar to the previous results, the differences in percent of peaks with footprints between target and background regions were smaller for the PWMs that are solely identified by HT (i.e., dark red dots labeled as ‘HT-only’ in **Supplementary Figure S6C**) when compared to PWMs identified by both methods. Furthermore, BiFET had higher enrichment p-values for these PWMs when compared to HT (**Supplementary Figure S6D**). Together these results suggest that footprint detection is subject to high rates of false positive calls and BiFET can be a useful downstream analysis method to reduce false positive associations for accurate interpretation of footprint enrichments.

### Background set choice affects false positive rate and detection power in standard tests

Simulation studies suggested that differences in read counts have a bigger impact on enrichment results than differences in GC content. Therefore, the differences between BiFET and HT enrichment results for PBMC-and islet-specific peaks likely stem from the differences in average read counts between target and background peaks (**Figures 3C, E**, left panel). To test this, we repeated HT enrichment analyses using different subsets of background peaks with different average read counts. First, we ordered background peaks based on their read counts and selected top n% of these peaks, where n is ranging from 50% to 100%, where 100% is equal to the original background set. Using these peak sets as the new background set, we performed HT and identified the set of PWMs significantly enriched in PBMC-specific peaks (Hn). As n increased from 50% to 100% (**Table 2A**), the average read counts of newly defined background peaks decreased from 168 to 93 and the number of identified PWMs (|Hn|) increased from 105 to 205. These analyses suggest that HT results highly depend on the choice of background regions and FP rate for enrichments increase as the difference between target and background regions increase in terms of their average ATAC-seq read counts. The PWMs captured by each of these analyses almost fully matched BIFET results (Hn ∩ B in Table 2A), suggesting again that BiFET captures likely true positives.

A potential solution to the dependence of HT results on background set choice is to carefully select background regions to match target regions in terms of GC content and average read counts. Sub sampling background regions to match the GC content of target and background regions has been widely used in motif enrichment analyses to correct for GC bias (19,20). However, in addition to the difficulty of sub-sampling background peaks to match target peaks both in terms of GC content and read counts simultaneously, there are several disadvantages associated with this strategy. First, having a smaller set of background peaks would reduce the power to detect differentially enriched PWMs. In our PBMC and islet data analyses, we had a large background set (n=40,116) and therefore sufficient power to detect enriched PWMs. Decrease in the size of background set can be tolerated up to a certain point. However, as background set shrinks further, the detection power would decrease. To test this, we randomly selected a subset of background peaks (n%) used in the most stringent case in the previous test (i.e., top 50% of the background peak). As n decreased from 100% (original set, 20,058 background peaks) to 10% (2005 background peaks), the number of enriched PWMs (i.e., |Hn|) also decreased from 105 to 33 (**Table 2B**), showing the reduction in power driven by the size of the background set. The second problem with random sampling of background peaks is the stochasticity it introduces in data analyses and the enrichment results. We tested this by repeating the random sampling of background peaks 10 times, where 10% of 20,058 peaks were selected as background peaks at each iteration. The number of PWMs significantly enriched in target peaks compared to these background peaks varied from 25 to 51 among different runs (**Table 2C**), with only 13 TFs common across 10 runs. These analyses suggest that the choice of background set has a significant impact on HT enrichment results and cannot be easily handled by subsampling data. BiFET does not require prior selection of background regions and works effectively with any background set, even if this set significantly differs from the target sets in terms of chromatin accessibility levels and GC content.

**Table 2.** HT results depend on background peaks used in the analyses. We tested different scenarios to understand the impact of background peak selection on HT results. (**A**) Selecting top n% of all background peaks based on their read counts showed that increasing the difference between target and background sets in terms of read counts increases the false positive enrichments with HT. (**B**) Randomly sampling different percentages of background peaks (n∼20,000 peaks) showed that reducing the size of the background set reduces detection power for HT. (**C**) Repeating the analyses from (B) for 10 times showed that random sampling introduces stochasticity in the HT enrichment results, where different sets of PWMs are captured to be enriched in each run.

A. Use top n% background peaks
| n | 50 | 60 | 70 | 80 | 90 | 100 |
| --- | --- | --- | --- | --- | --- | --- |
| mean read count | 168 | 147 | 129 | 114 | 102 | 93 |
| $\vert H_n\vert $ | 105 | 115 | 134 | 157 | 182 | 205 |
| $\vert H_n \cap B\vert $ | 87 | 88 | 88 | 89 | 89 | 89 |

B. Randomly select n% of X = top 50 % background peaks
| n% | 10 | 20 | 30 | 40 | 50 | 60 | 70 | 80 | 90 | 100 |
| --- | --- | --- | --- | --- | --- | --- | --- | --- | --- | --- |
| mean read count | 168 | 169 | 165 | 169 | 168 | 171 | 170 | 169 | 168 | 168 |
| $\vert H_n\vert $ | 33 | 57 | 69 | 90 | 84 | 92 | 96 | 100 | 105 | 105 |

C. Randomly select 10 % of X = top 50 % background peaks
| Run | 1 | 2 | 3 | 4 | 5 | 6 | 7 | 8 | 9 | 10 |
| --- | --- | --- | --- | --- | --- | --- | --- | --- | --- | --- |
| mean read count | 165 | 171 | 168 | 177 | 169 | 167 | 170 | 171 | 173 | 168 |
| $\vert H_{10}\vert $ | 25 | 43 | 34 | 49 | 39 | 34 | 51 | 43 | 26 | 45 |

### Footprints for high-GC motifs are captured in regions with high read counts

To understand the association between read counts and the footprint detection rate for each TF, we further studied the *α*_*k*_ parameter in our models. Higher values for *α*_*k*_ imply that the TF *k* can be detected in peaks with low read counts while lower values for *α*_*k*_ imply that the TF *k* can be detected mainly in peaks with high read counts (Methods). Using PIQ calls from PBMC and islet data, we identified TFs with high *α*_*k*_ values (>95^th^ percentile) and low *α*_*k*_ values (<5^th^ percentile) (**Supplementary Table S8**). We restricted our analysis to TFs that have footprints in at least 0.05% of all peaks (n=29 peaks), since the estimate *α*_*k*_ could be unstable for TFs with fewer footprints. As suggested by our model, TFs with low *α*_*k*_ (blue bars in **Figure 4A**, Supplementary **Figure S7A** for islet) were detected within peaks with high read counts (i.e., bigger peaks), whereas the TFs with high *α*_*k*_ (red bars in **Figure 4A**, **Supplementary Figure S7A** for islet) were detected within peaks with low read counts (i.e., smaller peaks). Surprisingly, we noted that the *α*_*k*_ estimates obtained from PBMC footprinting data were in agreement with those obtained from the islet footprinting data (Spearman correlation=0.88, **Figure 4B**), suggesting that the dependence of TF footprint detection rate on read counts (i.e., *α*_*k*_ parameter in our models) is specific to each TF and independent of the underlying cell type.

We did not detect a strong relationship between the length or the information content of a PWM and the corresponding TF’s *α*_*k*_ value (**Supplementary Figures S8A, B** for PBMC; **Supplementary Figures S9A, B** for islet). However, GC content of the PWMs (i.e., the average probability of having G or C within a motif) was inversely correlated with α_*k*_values (**Supplementary Figure S8C** for PBMC (p-value= 3.8e−13); **Supplementary Figure S9C** for islet (p-value=2.5e−12)), which implies that TFs with low *α*_*k*_ values i) tend to have high GC content PWMs and ii) are detected in regions that have high GC content. Indeed, regions that harbored footprints for low *α*_*k*_ TFs had higher GC content than regions harboring footprints for high *α*_*k*_ TFs (**Figure 4C** for PBMC; **Supplementary Figure S7B** for islet). This is likely due to the correlation between GC content and read counts (r=0.54, p-value<e−16; **Figure 4D** for PBMC; **Supplementary Figure S10** for islet and EndoC-ßH1), which might be related to the GC-specific cutting bias of Tn5 transposase (39) or PCR amplification bias towards GC-rich fragments (40). Due to this correlation between GC content and read counts, GC-rich motifs are more frequently detected in peaks with high read counts. Furthermore, since footprint detection rate is positively associated with number of reads, GC-rich motifs are more frequently detected in these analyses (**Supplementary Figure S8D** for PBMC; **Supplementary Figure S9D** for islet). However, we noted exceptions to this association between footprint detection rate and high GC and high read counts of genomic regions. For example, footprints of certain TFs (e.g. *TEAD1/3/4*) were detected within peaks with high read counts, but low-GC contents, suggesting they are more difficult to detect in open chromatin assays and require deeper sequencing.

**Fig. 4.**
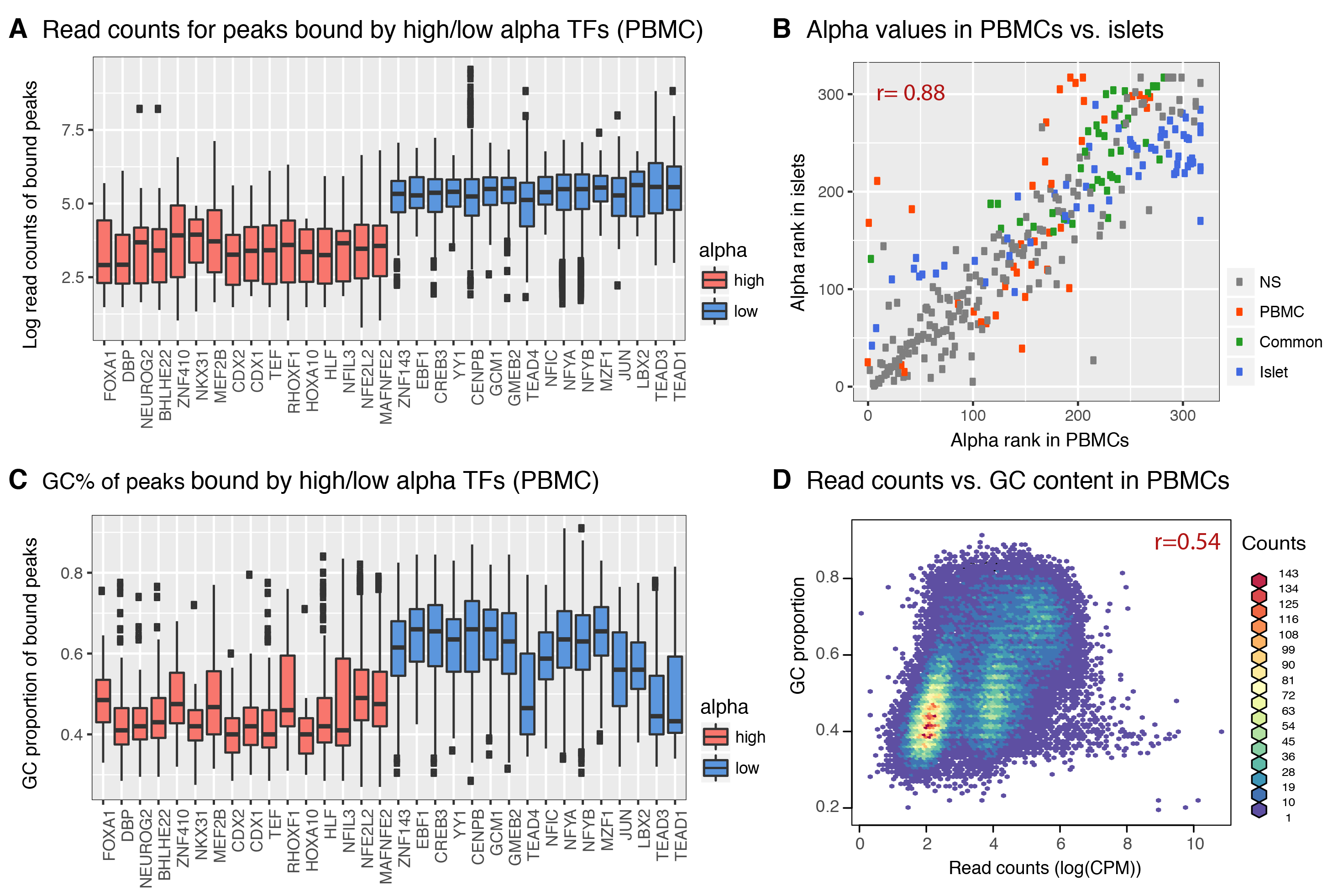
The relation between TF motif features and footprint detection rate. (**A**) Distribution of read counts for peaks that have footprints for high *α*_*k*_ TFs (above 95^th^ percentile of *α*_*k*_, red bars) and low values (below 5^th^ percentile, blue bars) in PBMCs. Footprints for low TFs were found in peaks with high read counts, whereas footprints for high *α*_*k*_ TFs were found in low read count peaks. (**B**) estimates obtained from PBMC footprint data correlate significantly with estimates from islet footprint data in rank. The TFs that are PBMC-specific are colored red, those that are islet-specific are colored blue, those that are both PBMC and islet-specific are colored green and those that are neither PBMC nor islet-specific are colored grey. (**C**) Distribution of GC contents for the peaks that have footprints for high TFs (above 95^th^ percentile, red bars) and low values (below 5^th^ percentile, blue bars) in PBMCs. (**D**) GC proportion of a region correlates significantly with the ATAC-seq read counts aligning to this location.

## DISCUSSION

In this study, we showed that TF footprint detection at a genomic locus is impacted by chromatin accessibility levels (i.e., ATAC-seq read count) and the GC content of this genomic region. This dependence is critical and needs to be taken into consideration in enrichment analyses while comparing target regions to background regions. For this purpose, we developed BiFET, a novel enrichment test that corrects for the differences in sequence and read counts of target and background regions. We applied BiFET on ATAC-seq data from the human beta cell line EndoC-ßH1 using TF footprints called with CENTIPEDE (6), HINT-BC (25) and PIQ (7) as well as on ATAC-seq data from human PBMCs and islets to demonstrate that BiFET can effectively identify potential regulators of cell-type specific loci.

Our simulation results showed that BiFET is a robust alternative to standard enrichment tests, e.g., hypergeometric test (**Table 1**). For footprinting data analyses, standard tests are very sensitive to the choice of background regions and require these regions to be comparable to target regions in terms of average read counts and GC content. If the background regions are not properly selected in such analyses, which has its own challenges (**Table 2**), they lead to high false positive rates and therefore spurious associations between open chromatin regions and TFs. BiFET on the other hand does not require selecting background regions as it accounts for any differences between target and background loci in terms of GC content and read counts. Overall, BiFET reduces false positive rates and provides a high detection power. Furthermore, we noted similar improvements in enrichment analyses using BIFET with footprints called via three different methods (CENTIPEDE, HINT-BC, and PIQ), suggesting that BiFET works effectively regardless of the algorithm used for calling TF footprints.

The distribution of read counts across the genome is confounded with the cleavage bias of cutting enzymes used in chromatin accessibility assays (4,21). For example, Tn5 transposase used in ATAC-seq libraries is biased towards more frequently cutting guanosine-and cytidine-rich sequences, thus, regions with high GC content tend to have more cleavages in such assays (39), however very little is known about the impact of this bias on TF footprinting data analyses (22). In agreement with the reported sequence biases, we observed that read counts and GC contents were positively associated in all ATAC-seq datasets studied here regardless of the cell types. Furthermore, we observed that TFs with GC-rich motifs are detected more frequently in regions with higher read counts, which also typically have high GC contents. This observation further supports that it is necessary to adjust for the potential biases in the data in TF footprint enrichment analysis.

Although TF footprinting provides an attractive and cost-effective alternative to ChIP-seq assays, it is prone to false positive calls as also suggested by our analyses using the randomly generated motifs. Therefore, an enrichment test that can reduce false positive associations between TFs and genomic regions is critical to effectively analyze and interpret TF footprinting data. Another pitfall of TF footprinting analysis is the high false negative detection rate. It is known that some TFs leave no footprints despite prominent binding to DNA (8,41). Furthermore, we observed that some TFs with known cell-specific functions were missed in the enrichment test due to i) missing PWMs or ii) small numbers of footprints detected for these TFs such as PDX1 and NKX6−1 for islets, which both have AT-rich PWMs. These are some of the open challenges that still hinder footprinting analyses, which ongoing studies are trying to address (42,43).

In summary, we observed that there is a positive association between read counts and GC content of a given locus and the number of TF footprints detected at this site. If not taken into consideration, this association significantly inflates the false positive rate in enrichment tests. By modeling this association and accounting for this bias, BiFET reduces false positive rate without compromising the true positive rate. This advanced and novel test is more effective for the analyses and interpretation of TF footprinting data that is inherent to biases and can distinguish the most probable regulators of cell-or disease-specific functions from potentially spurious ones, which will be an essential next step in genomic medicine studies that are generating chromatin accessibility maps from clinically-relevant samples to study complex human diseases (10–13).

## AVAILABILITY

‘BiFET’ and all associated source code is freely available as a Bioconductor package and at our GitHub page: https://github.com/UcarLab/BiFET.

## SUPPLEMENTARY DATA

Supplementary Data are available at NAR Online.

## FUNDING

This study was made possible by generous financial support of the National Institute of General Medical Sciences (NIGMS) under award number GM124922 (to DU) and by The Jackson Laboratory startup funds (to DU). Opinions, interpretations, conclusions, and recommendations are solely the responsibility of the authors and do not necessarily represent the official views of the National Institutes of Health (NIH).

## CONFLICT OF INTEREST

None declared.

